# Programmed mutation of liver fluke granulin using CRISPR/Cas9 attenuates virulence of infection-induced hepatobiliary morbidity

**DOI:** 10.1101/386219

**Authors:** Patpicha Arunsan, Wannaporn Ittiprasert, Michael J. Smout, Christina J. Cochran, Victoria H. Mann, Sujittra Chaiyadet, Shannon E. Karinshak, Banchob Sripa, Neil D. Young, Javier Sotillo, Alex Loukas, Paul J. Brindley, Thewarach Laha

## Abstract

Infections with several flatworm parasites represent group 1 biological carcinogens, *i.e.* definite causes of cancer. Infection with the food-borne liver fluke *Opisthorchis viverrini* causes cholangiocarcinoma (CCA). Whereas the causative agent for most cancers, including CCA in the West, remains obscure, the principal risk factor for CCA in Thailand is opisthorchiasis. We exploited this established link to explore the role of the secreted parasite growth factor termed liver fluke granulin (*Ov*-GRN-1) in pre-malignant lesions of the biliary tract. We targeted the *Ov*-*grn-1* gene for programmed knockout and investigated gene-edited parasites *in vitro* and in experimentally infected hamsters. Both adult and juvenile stages of the liver fluke were transfected with a plasmid encoding a guide RNA sequence specific for exon 1 of *Ov-grn-1* and the Cas9 nuclease. Deep sequencing of amplicon libraries from genomic DNA from gene-edited parasites exhibited programmed, Cas9-catalyzed mutations within the *Ov-grn-1* locus, and tandem analyses by RT-PCR and western blot revealed rapid depletion of *Ov-grn-1* transcripts and protein. Newly excysted juvenile flukes that had undergone editing of *Ov-grn-1* colonized the biliary tract, grew and developed over a period of 60 days, were active and motile, and induced a clinically relevant pathophysiological tissue phenotype of attenuated biliary hyperplasia and fibrosis in comparison to infection with wild type flukes. This is the first report of gene knock-out using CRISPR/Cas9 in a parasitic flatworm, demonstrating the activity and utility of the process for functional genomics in these pathogens. The striking clinical phenotype highlights the role in virulence that liver fluke growth factors play in biliary tract morbidity during chronic opisthorchiasis.

## Introduction

Liver fluke infection caused by species of *Opisthorchis* and *Clonorchis* remains a major public health problem in East Asia and Eastern Europe. *O. viverrini* is endemic in Thailand and Laos, where ∼10 million people are infected with the parasite (*1*). There is no stronger link between a human malignancy and a parasitic infection than that between CCA and infection with *O. viverrini* (*2*). In endemic regions such as Northeastern Thailand, infection causes hepatobiliary diseases including cholangitis and periductal fibrosis - a major risk factor for CCA (*3*). The north of Thailand suffers the highest incidence of CCA in the world, often exceeding 80 cases per 100,000 population, and for which up to 20,000 people annually are admitted for surgery. Unfortunately, prognosis for fluke-induced cancer is poor (*1, 4, 5*).

How and why opisthorchiasis induces cholangiocarcinogenesis is likely multi-factorial, including mechanical irritation of the biliary tract during migration and feeding of the liver fluke, inflammatory molecules released by the parasite, and nitrosamines in fermented foods that are a dietary staple in northern Thailand. To survive in the hostile host environment, parasitic helminths produce an assortment of excretory/secretory (ES) products including proteins with diverse roles at the host–parasite interface. This interaction has long been thought but not fully understood to modify cellular homeostasis and contribute to malignant transformation during chronic opisthorchiasis (*6*). Feeding activity of the liver fluke inflicts wounds in the biliary tree, lesions that undergo protracted cycles of repair and re-injury during chronic infection. The liver fluke secretes mediators that accelerate wound resolution in cholangiocytes, an outcome that can be compromised following silencing of expression of *Ov-grn-1* using RNA interference (*7, 8*). We hypothesize that proliferation of biliary epithelial cells induced by *Ov*-GRN-1 is a pivotal factor in establishing a tumorigenic microenvironment in livers of infected individuals.

Progress with development of genetic tools for functional genomic studies with platyhelminth parasites has been limited to date (*9*). The use of clustered regularly interspaced short palindromic repeats (CRISPR) associated with Cas9, an RNA-guided DNA endonuclease enzyme, has revolutionized genome editing in biomedicine, agriculture and biology at large (*10, 11*). Progress with CRISPR/Cas9 in numerous eukaryotes including the nematodes *Caenorhabditis elegans* and *Strongyloides stercoralis* has been described (*11-14*), but this form of gene editing has not been reported for flatworm parasites. Here, we deployed CRISPR/Cas9 to knockout (mutate) the *Ov-grn-1* gene and assess the virulence of gene-edited flukes *in vitro* and *in vivo* in a hamster model of opisthorchiasis.

## Results and Discussion

### Growth factor secreted by carcinogenic liver mutated by programmed genome ediitng

A construct termed pCas 9-*Ov-grn-1* (Fig. 1a and Supplementary Fig. 1a) was assembled following assessment of the *Ov-grn-1* locus in the annotated genome sequence of *O. viverrini* (*15*). Nucleotide sequences in pCas 9-*Ov-grn-1* were confirmed by Sanger cycle sequencing; pCas 9-*Ov-grn-1* encodes the Cas9 nuclease and a single guide RNA (sgRNA) complementary to a target sequence 5’-GATTCATCTACAAGTGTTGA within exon 1 of *Ov-grn-1*. The predicted programmed cleavage site was predicted to be at three residues upstream of a CGG proto-spacer adjacent motif sequence (PAM) in exon 1 of *Ov-grn-1* (Fig. 1b; Supplementary Fig. 1b). Adult *O. viverrini* flukes recovered from experimentally infected hamsters were subjected *in vitro* to square wave electroporation (*8*) in the presence of pCas 9-*Ov-grn-1* DNA, and thereafter maintained in culture for three weeks. The activity of CRISPR/Cas9 was evaluated by two approaches.

**Figure 1.**
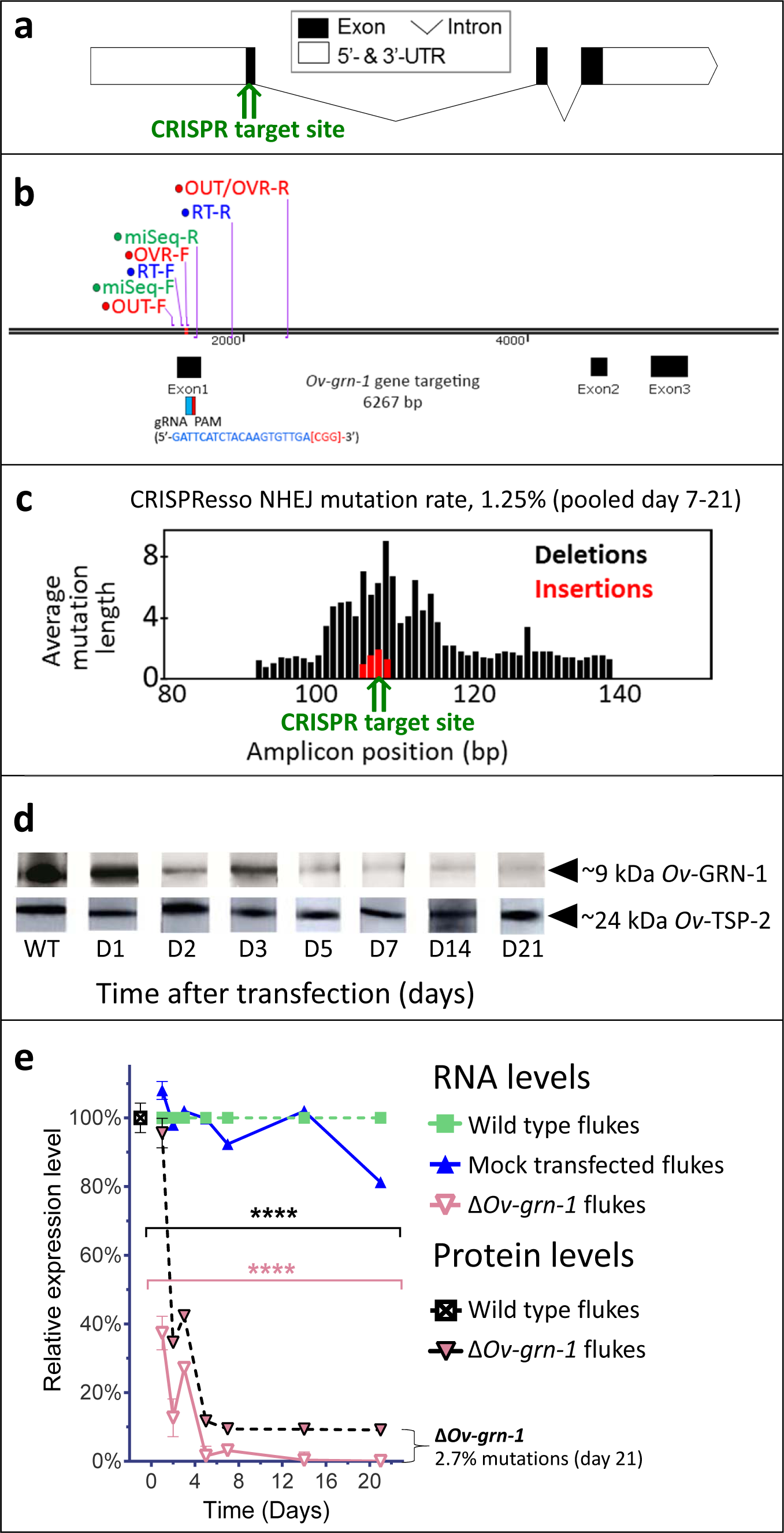
CRISPR/Cas9-mediated gene editing strategy to knock-out *Ov-grn-1* in adult *Opisthorchis viverrini* liver flukes. **a**, Schematic depiction of *Ov-grn-1* gene with CRISPR/Cas9 target site in first exon marked with green arrow. **b**, The exon1, 2 and 3 location of *Ov-grn-1* gene, size of 6,267 bp and gRNA targeting exon1: gRNA sequence is blue color and PAM is red in bracket. Primer pairs were used to detect mRNA expression levels by using RT-qPCR assay (*Ov-grn-1* RT- forward or RT-F and *Ov-grn-1* RT-reverse or RT-R), Tri-primers were used to detect the % relative fold amplicon or mutations (outside-forward or OUT-F, overlap-forward or OVR-F and reverse primer or OUT/OVR-R) and MiSeq forward and reverse (MiSeq-F and MiSeq-R) primers were used to prepare the NGS amplicon. **c**, CRISPR/Cas9-induced adult fluke genomic insertion (red bars) and deletion (black bars) mutations (INDELs) detected in the *Ov-grn-1* gene; CRISPR target site is denoted by the green arrow. Average mutation length is plotted against *Ov-grn-1* gene amplicon position in base pairs (bp). **d**, Somatic tissues of individual adult worms (in triplicate per time per group) were solubilized, electrophoresed in SDS-PAGE gels, transferred to nitrocellulose membrane and probed with anti-*Ov-*GRN-1 rabbit antibody. WT: wild type control fluke tissues; D1-21: Δ*Ov-grn-1* fluke tissues sampled the arrow highlighting the ∼9 kDa *Ov-*GRN-1 band at different days post-transfection and Δ*Ov-grn-1* flukes show similar *Ov-*TSP-2 protein (control antibody) expression levels. D1-21 = protein products from flukes day 1-21 post Δ*Ov-grn-1* treatment. Western blot panels of probed with anti-*Ov-*TSP-2 rabbit antibody, the arrow highlighting the ∼24 kDa *Ov-*TSP-2 band. **e**, Reduced expression of *Ov-grn-1* mRNA and *Ov-*GRN-1 protein after transfection of adult flukes with *Ov-grn-1* CRISPR/Cas9 construct using quantitative real-time PCR (mRNA) and densitometry conversion of Western blot signals (protein). Data are plotted relative to wild type (WT) fluke values (100%) as an average percentage from 3 replicates with SD error bars. **** = *P*<0.0001 compared to WT fluke protein (black) or RNA (pink) at each time point with two-way ANOVA Holm-Sidak multiple comparison test.

First, quantitative PCR (qPCR) was employed, which relies on the inefficiency of binding of a primer (here termed OVR-F) overlapping the target genomic sequence of the gRNA, i.e. where mutations are expected to have occurred, compared to the binding efficiency of flanking primers, i.e. outside the mutated region (*16, 17*) (flanking primers termed OUT and OUT-R) (Fig. 1b). Genomic DNA (gDNA) templates were investigated by quantitative PCR to quantify the efficiency of programmed gene-editing at the target locus; the ratio between the OVR-OUT-R products and OUT-F-OUT-R products provided an estimate of the amplification fold-reduction in the sample of CRISPR/Cas9-edited compared to gDNA from control, wild type liver flukes at the target sequence of the sgRNA, i.e. the annealing site for the OVR primer. A reduction in relative fold amplification of 2.7% was detected in gDNA from the Cas9-treated worms (Fig. 1e; Supplementary Fig. 1c).

Second, to identify and quantify the mutations that arose in the genome of *Ov-grn-1*-edited (termed Δ*Ov-grn-1*) flukes, we used an amplicon-sequencing approach. A targeted (amplicon) sequence library was constructed from genomic DNA from some of the flukes. A 173 bp fragment spanning the predicted double stranded break site of *Ov-grn-1* was amplified from the gDNAs primed with oligonucleotides flanking 1496-1668 nt of *Ov-grn-1.* Adaptors and barcodes were ligated into these amplicon libraries. The MiSeq libraries were undertaken by Illumina MiSeq-based deep sequencing. Insertion-deletion (INDEL)/mutation profiles in the sequence reads were compared in multiple sequence alignment with wild type reference template sequence (1496-1668 nt) of *Ov-grn-1*. Each amplicon was sequenced on the MiSeq Illumina platform and quantified the gene editing frequency by the CRISPResso pipeline (*18, 19*). More than 2 million sequenced reads were aligned against the reference sequence, which predicted the presence of 27, 616 non-homologous end joining (NHEJ) reads, specifically 170 reads with insertions (0.6%), 193 reads with deletions (0.7%) and 27,277 reads with substitutions (98.7%). At large, 1.25% of the sequenced reads exhibited NHEJ mutations (Fig. 1c). Among these NHEJ reads, there were >100 forms exhibiting mutations that would disrupt the coding sequencing of *Ov-grn-1.* Four representatives of the INDEL-bearing traces aligned with the WT allele are presented in Supplementary Fig. 1b. The Illumina sequencing reads are available as GenBank accessions SRR5764463-5764618, at https://www.ncbi.nlm.nih.gov/Traces/study/?acc=SRP110673, Bioproject, www.ncbi.nlm.nih.gov/bioproject/PRJNA385864.

Effects of gene editing on transcription and protein expression were investigated. Levels of both *Ov-grn-1* mRNA transcripts as determined by RT-PCR and of granulin, as detected by western blot with anti-*Ov*-GRN-1 sera, fell significantly from days 1 and 2 after transfection, respectively (*P* ≤ 0.0001; Fig. 1d, e; Supplementary Fig. 1c). Expression levels of two reference genes, the actin transcript (Supplementary Fig. 1c) and the *Ov-*TSP-2 tegument protein (Fig. 1d), were not influenced by the programmed mutation of *Ov-grn-1*.

### Mutaion of liver fluke granulin attenuates infection-induced biliary tract morbidity

To investigate whether gene editing of *Ov-grn-1* impacted *in vitro* indicators of pathogenesis, the capacity of ES products from WT, mock-transfected and gene-edited flukes to drive proliferation and scratch wound repair of the H69 human primary cholangiocyte cell line was assessed. ES from WT and mock-transfected adult flukes stimulated cell proliferation and wound closure whereas an equivalent amount of ES products from Δ*Ov-grn-1* flukes resulted in significantly reduced cell proliferation over the six day course of the assay (*P* ≤ 0.0001; Fig. 2a, b; Supplementary Fig. 2a, b) and significantly reduced *in vitro* wound closure over 36 hours (*P* ≤ 0.0001; Fig. 2c; Supplementary Fig. 2c, d), consistent with a reduction in the amount of *Ov*-GRN-1 secreted from the gene-edited liver flukes.

**Figure 2.**
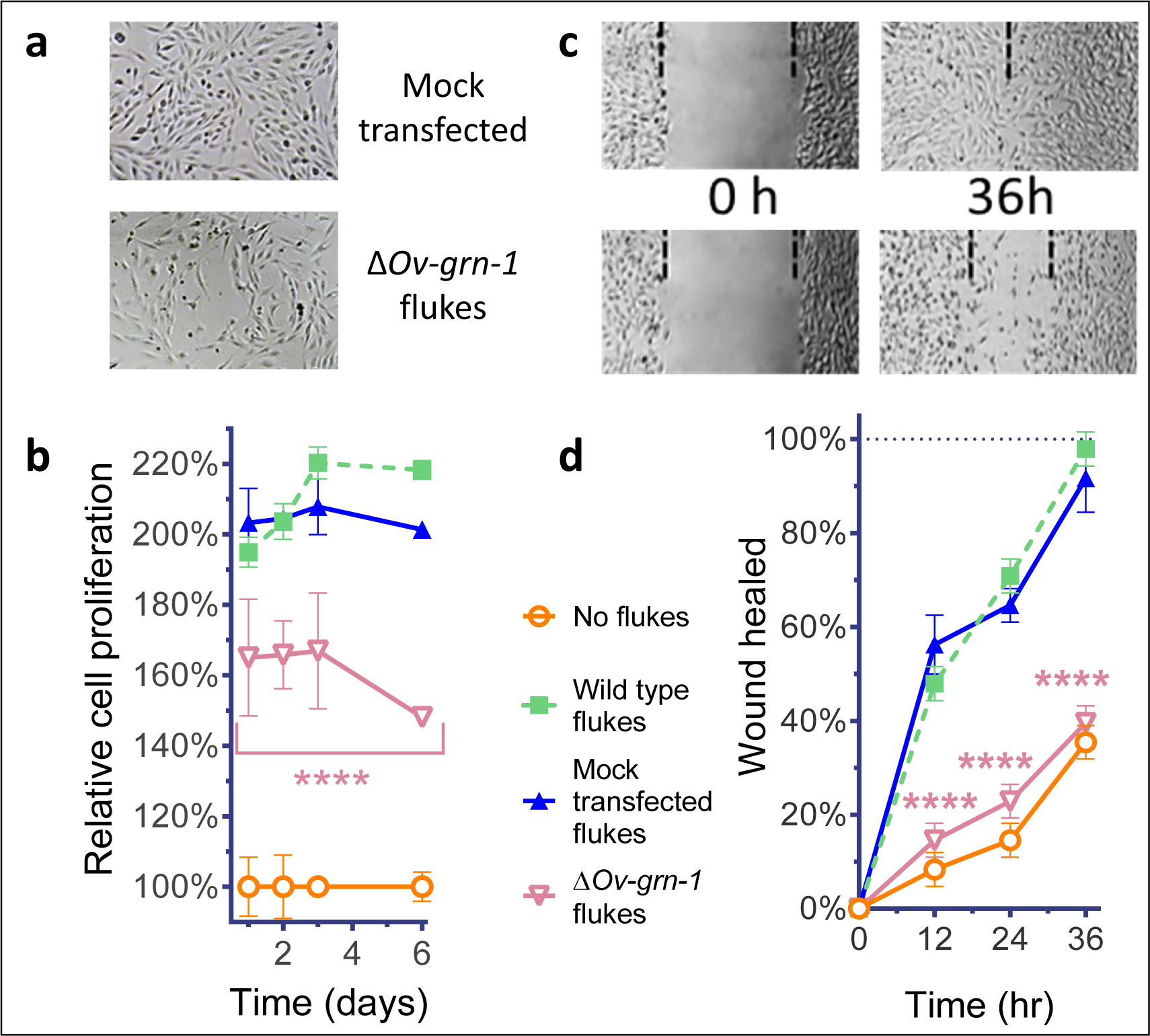
Δ*Ov-grn-1* adult fluke ES products induce less *in vitro* cell proliferation and wound repair. **a**, Representative cell proliferation images of H69 immortalized human cholangiocyte cell line co-cultured with flukes in Trans-well plates; mock transfected (top) and Δ*Ov-grn-1* (bottom) groups are shown at day 3. **b**, Reduced cell proliferation induced by *ΔOv-grn-1* fluke ES products, as shown in panel a, quantified from days 1-6. Data is plotted as average relative percentage to cells cultured with “no flukes”. **c**, Representative image of H69 cholangiocyte scratch wound repair when cells were co-cultured in Trans-well plates with flukes. Mock transfected (top) and Δ*Ov-grn-1* (bottom) groups are shown at 0 and 36 h post-scratch wounding. Dotted line shows the edge of the wound. **d**, Scratch wound repair assay quantified from 0 to 36 hours, reveling diminished healing in the Δ*Ov-grn-1* group. Panels **b** and **d**: mean ± SD, three replicates; \*\*\*\**P*<0.0001 compared to wild type flukes with two-way ANOVA Holm-Sidak multiple comparison test.

Notwithstanding the noteworthy effects observed with gene-edited, adult developmental forms, the metacercaria (MC) (Fig. 3a) is the infective stage of *O. viverrini* for humans. Accordingly, we investigated gene knockout in MC. No effect was apparent on *Ov-grn-1* transcript levels in MC, (Supplementary Fig. 3), suggesting that delivery of the pCas 9-*Ov-grn-1* by electroporation through the MC cyst wall was ineffective. Exposure to bile acids and gastric enzymes results in excystation of *O. viverrini* MC in the duodenum of the infected mammalian host. We mimicked this process *in vitro* using trypsin to release the newly excysted juvenile worms (NEJ; Fig. 3b), after which NEJ were subjected to electroporation with CRISPR/Cas9 constructs as described for adult flukes. Marked depletion of *Ov-grn-1* transcripts (*P* < 0.0001) followed this manipulation (Fig. 3c).

**Figure 3.**
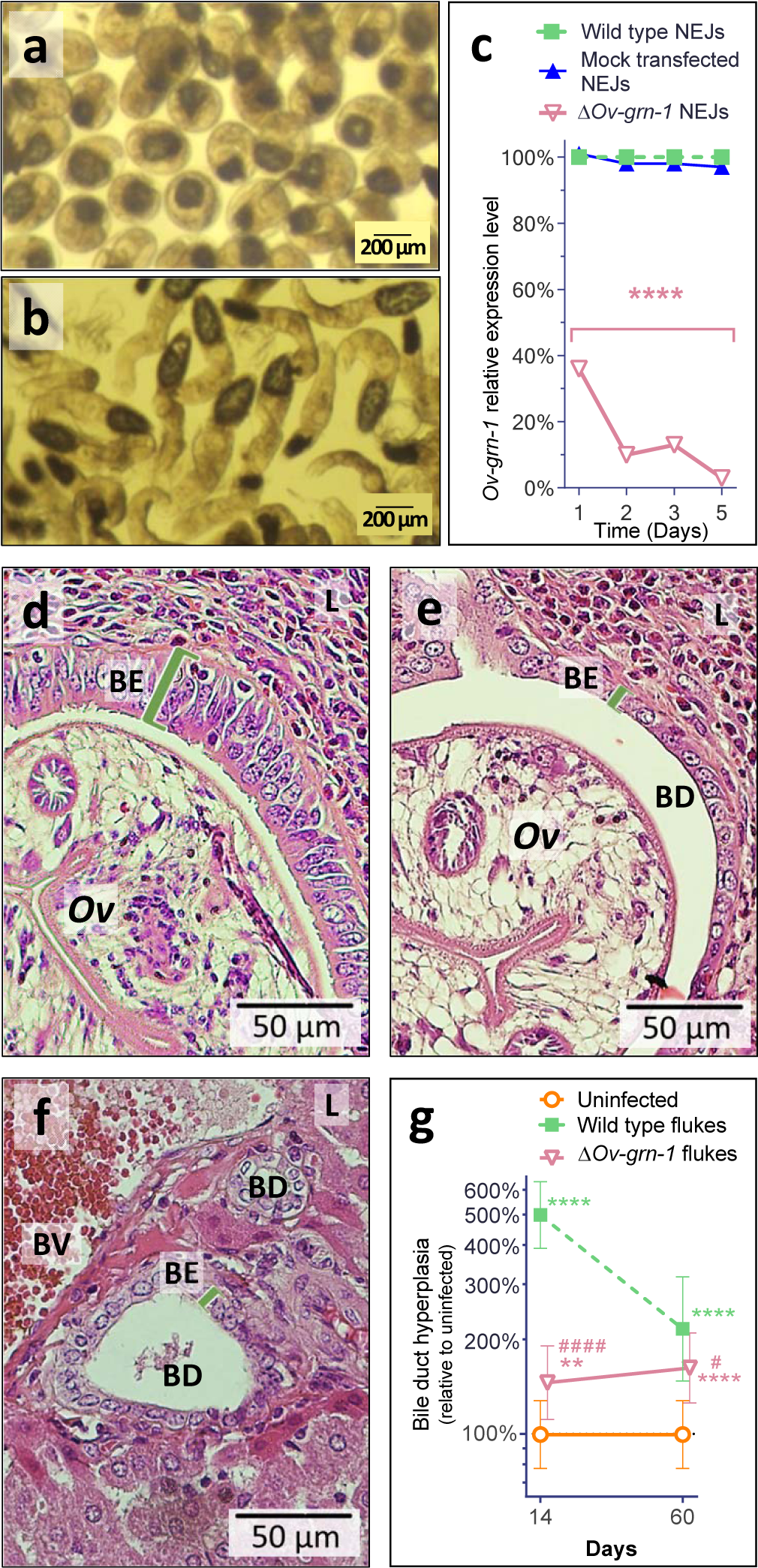
Δ*Ov-grn-1* newly excysted juveniles can infect hamsters and drive reduced short-term pathology. Images of infective metacercariae (**a**) and newly excysted juvenile flukes (NEJ) (**b**). **c**, *Ov-grn-1* mRNA expression in mock-transfected and Δ*Ov-grn-1* flukes over 5 days as quantified with qPCR and plotted relative to the wild type (WT) untreated group; mean ± SD, three replicates; **** = *P*<0.0001 compared to WT flukes with two-way ANOVA Holm-Sidak multiple comparison test at each time point. **d**, Representative image of 400x magnification of H&E stained thin section from hamster liver at 14 days after infection with WT flukes. **e**, Representative image of H&E stained thin sections showing hamster liver 14 days after infection with Δ*Ov-grn-1* flukes. **f**, Representative image of H&E stained thin section of control, uninfected hamster liver, which shows healthy, organized pavement-like profile of the cells of the biliary epithelium (BE) enclosing the lumen of the bile duct (BD) near a blood vessel (BV) within the liver (L). Infection by WT flukes (panel **d**) revealed thickened, disordered epithelium adjacent to the parasite (*Ov*). Infection with Δ*Ov-grn-1* flukes (panel **e**) revealed a bile duct epithelium more closely resembling the normal, uninfected hamster. **g**, Epithelium width/hyperplasia (green bracket) was quantified using ImageJ and plotted as the mean ± SD of five biological replicates (hamsters). Significant difference were apparent when compared to the uninfected group using the two-way ANOVA with Holm-Sidak multiple comparison test: \*\**P*<0.01 and \*\*\*\**P*<0.0001, and wild-type compared to Δ*Ov-grn-1*, ^#^ *P*<0.05 and ^####^*P*<0.0001.

In parallel, hamsters were infected, using gastric gavage, with 100 Δ*Ov-grn-1* NEJ or WT NEJ immediately after electroporation. At necropsy of the hamsters two to three weeks later, it was clear that Δ*Ov-grn-1* flukes had colonized the bile ducts in similar numbers to WT flukes, and were similarly motile. Strikingly, the infection with Δ*Ov-grn-1* parasites failed to induce the marked hyperplasia of the bile duct epithelia characteristic of infection with WT flukes. Specifically, infection with WT flukes had induced markedly disordered, hyperplasic growth of the epithelium adjacent to the parasites; at day 14, five times more than in uninfected controls (*P*≤ 0.0001) (Fig. 3d) whereas infection with the Δ*Ov-grn-1* flukes (Fig. 3e) provoked 11-fold less (*P* ≤ 0.0001) biliary hyperplasia than WT fluke infected livers (day 14, 145% thickening compared to uninfected controls; *P* ≤ 0.01) (Fig. 3g). Indeed, the bile ducts from hamsters infected with the Δ*Ov-grn-1* flukes generally resembled those of the control, uninfected hamsters (Fig. 3f).

To assess long-term survival of Δ*Ov-grn-1* NEJ in hamsters and associated chronic biliary morbidity, hamsters were infected with Δ*Ov-grn-1* and WT NEJ, and adult flukes were recovered and counted from the livers 60 days post-infection. Similar numbers of worms were recovered from both control and gene-edited liver fluke-infected hamsters (Fig. 4a). To assess the impact of infection with Δ*Ov-grn-1* on markers of chronic opisthorchiasis such as biliary fibrosis, liver sections from infected hamsters were stained with Picro-Sirius Red to detect collagen bundles in the biliary tract (Fig. 4b). Minimal deposits of collagen were seen in the periductal regions of the biliary tract of the control, uninfected hamsters whereas hamsters infected with WT flukes had thick bands of collagen surrounding enlarged bile ducts in the vicinity of the flukes. Significantly less collagen (28%) was detected in periductal sites of the biliary tract infected with Δ*Ov-grn-1* flukes (*P* < 0.001) compared to livers of hamsters infected with WT flukes (Fig. 4b, c). To further assess fibrosis in hepatobiliary tract, we stained for smooth muscle actin (ACTA2), an established marker of hepatic fibrosis (*20*), by probing thin tissue sections with anti-ACTA2 antibody. Livers of hamsters infected with WT flukes showed densely packed collagen fibrils that stained for ACTA-2 in periductal regions proximal to the parasites. In contrast, livers from hamsters infected with Δ*Ov-grn-1* flukes displayed an irregular distribution of less dense collagen fibrils with less ACTA2-specific fluorescence (Fig. 4d; Supplementary Fig. 4). The intensity of ACTA2-specific fluorescence was quantified by measuring fluorescence intensity; livers from Δ*Ov-grn-1* fluke-infected hamsters showed 94% reduction (*P* <0.01) of median fluorescence values relative to controls infected with WT flukes (Fig. 4e).

**Figure 4.**
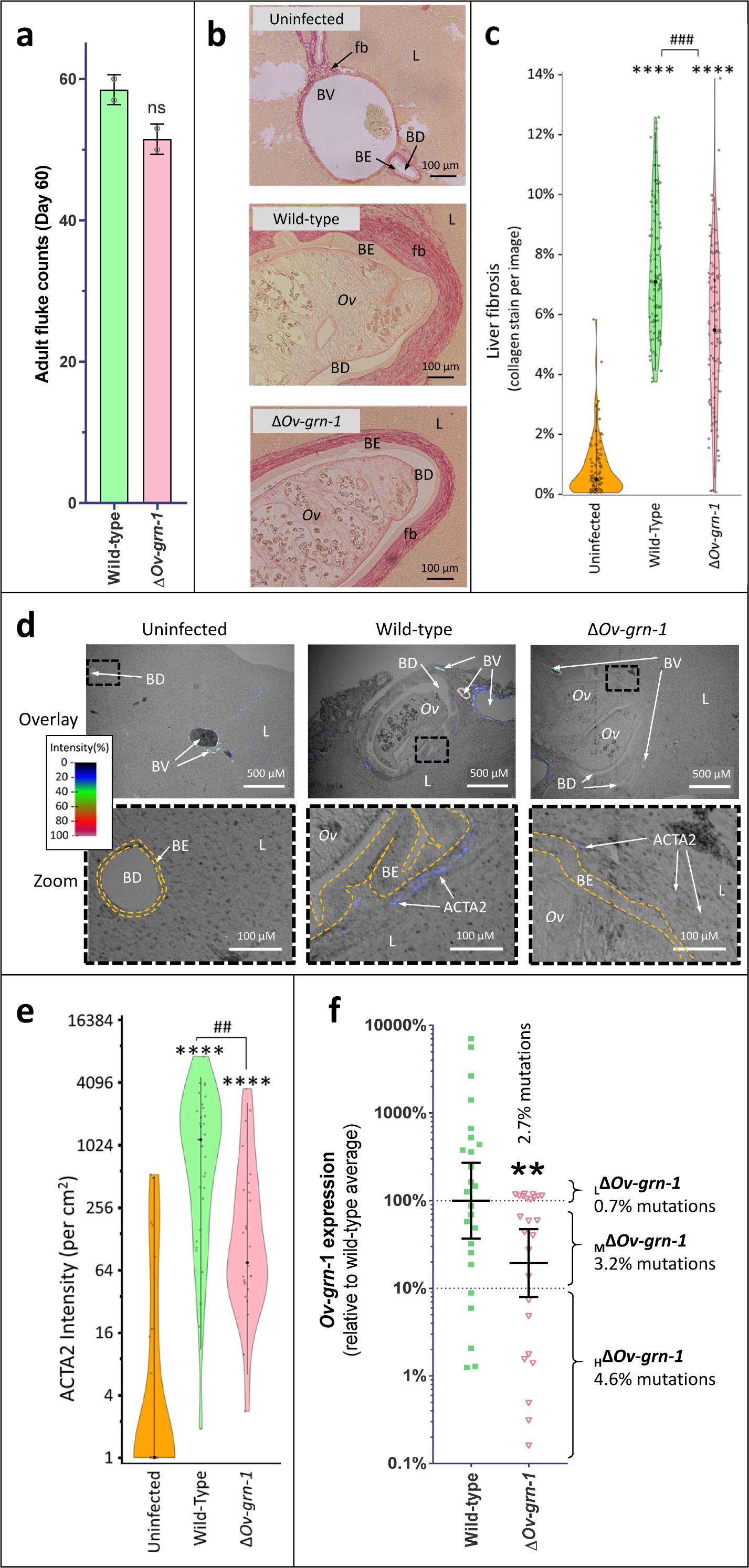
Diminished level of fibrosis in hamsters during chronic infection with gene-edited Δ*Ov-grn-1* liver flukes. Panel **a**, Adult fluke numbers were counted from the livers at day 60 post-infection and shown as the average and range for two hamsters per group. Fluke counts showed minimal (12%; *ns*) difference between the wild type (WT) and Δ*Ov-grn-1* groups. **b,** Representative images of Sirius red stained hamster liver (L) sections from uninfected animals show minimal collagen fibrosis (fb) deposition, stained red surrounding the blood vessel (BV) and bile ducts (BD) with normal biliary epithelia (BE). Livers from hamsters infected with WT flukes (*Ov*) show heavy collagen deposition with elongated BE cells adjacent to resident flukes.. Δ*Ov-grn-1* fluke infected livers showed substantial collagen deposition compared to uninfected liver sections, yet showed far less than the livers infected with WT flukes. **c**, Liver fibrosis quantified with ImageJ MRI-fibrosis plugin shown as a violin plot. 100 images containing bile ducts from 20 sections (5 animals) per group showing SD as a vertical line with the median indicated by the central black dot. Δ*Ov-grn-1* group showed a 23% reduction in median fibrosis compared to WT group. The width of the “violin-shape” represents measurement frequency. Kruskal-Wallis with Dunn’s multiple comparison test used to compare groups. Against uninfected: \*\*\*\**P*<0.0001, and Δ*Ov-grn-1* against WT: ### *P*<0.001. **d**, Representative immunofluorescence/bright field overlay images of liver sections probed with anti-ACTA2 antibody with fluorescence intensity shown on a blue/green/red scale. ACTA2 protein is always detected in my fibroblasts surrounding blood vessels, but not near normal bile ducts. When detected near the BE layer it is suggestive of myofibroblast generation as a response to BE damage. The upper panels (Overlay) show a wide view of the liver sections with a boxed region that highlights a section of interest that is magnified in the lower panels (Zoom). Hamster liver sections exhibited intense fluorescence (arteries: red/green, veins: blue/green) surrounding BVs whereas only minimal fluorescence was seen near BD in livers of uninfected hamsters The highlighted magnified (Zoom) panels show WT infected animal livers with weak (blue) but consistent ACTA2 staining surrounding thickened BE layer (inner and outer cell edge marked with orange dotted line) around BDs with WT flukes. Livers from hamsters infected with Δ*Ov-grn-1* flukes show weak and irregular ACTA2 staining surrounding BD. **e**, Quantified levels of ACTA2 surrounding BDs from hamster liver sections. Violin plot with reverse log2 Y-axis showing the ACTA2 intensity (per cm^2^ at 300 PPI) surrounding BE from 25-30 discrete BD images per group (three hamsters) assessed with ImageJ. Zero values from uninfected group were deemed to have a value of 1 so that a log axis could be plotted. SD indicated as a line with mean value marked as a central black dot and the violin-shape width representing measurement frequency. ACTA2 staining shows 94% median reduction in Δ*Ov-grn-1* fluke-infected livers compared to WT fluke-infected animals. One-way ANOVA with Holm-Sidak multiple comparison test, ****P<0.0001 compared to uninfected and ##P<0.001 compared to Δ*Ov-grn-1* flukes against WT flukes. **f**, *Ov-grn-1* mRNA expression levels are reduced in Δ*Ov-grn-1* flukes compared to WT flukes. Reverse log10 Y-axis shows the qPCR 2^(-ΔΔCt) results from flukes 60 days after programmed CRISPR/Cas9 gene editing and hamster infection plotted relative to mean value of the WT infection. The WT group showed a broad level of expression; the mean expression level for the Δ*Ov-grn-1* flukes was 19.4% of the WT group; Mann-Whitney nonparametric test, \*\**P*<0.01. While significantly reduced overall, the Δ*Ov-grn-1* fluke group ranged from no apparent effect to markedly diminished expression of *Ov-grn-1*. Mutation frequency was assessed by assigning the Δ*Ov-grn-1* group of flukes into three sub-groups based on CRISPR/Cas9 mutation frequency. Eight flukes with highly effective CRISPR gene knock-out (Δ*Ov-grn-1*: *Ov-grn-1*<10% expression), seven flukes with modest levels of transcript knock-out (Δ*Ov-grn-1*: 10-70% *Ov-grn-1*) and 10 flukes with little or no effect (Δ*Ov-grn-1*: *Ov-grn-1* 100-120%). The mean mutation frequency amongst the three Δ*Ov-grn-1* sub-groups was 2.7%.

Despite reaching high levels of significance, the data range was high for both markers of collagen deposition assessed. We hypothesized that this was likely due to inconsistencies in the electroporation-mediated delivery of the CRISPR/Cas9 constructs between individual flukes. We therefore collected Δ*Ov-grn-1* and WT flukes from hamster bile ducts 60 days post-infection and assessed *Ov-grn-1* gene expression from individual flukes (Fig. 4f). We then separated flukes into three groups based on *Ov-grn-1* mRNA expression levels: >100% relative to WT average; 10-100% relative to WT average; <10% relative to WT average. We pooled genomic DNA from flukes to form the three groups described above and assessed mutation frequencies. In line with the *Ov-grn-1* mRNA expression profiles, Δ*Ov-grn-1* flukes where gene editing appeared to be relatively inefficient (i.e. >100% mRNA expression relative to WT average) had an average mutation frequency of just 0.7%, whereas Δ*Ov-grn-1* flukes where gene editing appeared to be moderately efficient (10-100% mRNA expression relative to WT average) had an average mutation frequency of 3.2%; Δ*Ov-grn-1* flukes where gene editing appeared to be highly efficient (<10% mRNA expression) had an average mutation frequency of 4.6% (Fig. 4f). The combined mutation frequency was 2.7% (Fig. 4f), notably similar to the 2.7% mutation rate detected when Δ*Ov-grn-1* flukes were cultured *in vitro* for 21 days (Fig 1e and Supplementary Fig. 1c).

In overview, we describe the first example of successful somatic gene editing of a parasitic flatworm using CRISPR/Cas9. The results revealed that gene editing induced disruption of expression of *Ov-grn-1* in liver flukes, which in turn revealed a pathologically relevant phenotype in the mammalian biliary tract. Following programmed gene editing, the lesion was apparently repaired by non-homologous end joining. The bacterial Type II Cas9 system is active in this liver fluke, and we conjecture that Cas9-catalyzed gene editing will be active in trematodes and parasitic platyhelminths at large. As noted, whereas the causative agent for many cancers, including CCA in the West, remains obscure, the principal risk factor for CCA in Thailand and Laos has long been established - infection with *O. viverrini*. Cas9-based gene editing and the hamster model of human opisthorchiasis utilized herein, including genetic manipulation of the infective NEJ parasite, together provide a facile, functional genomics system to interrogate parasite pathogenicity and carcinogenicity in an informative rodent model of liver fluke infection-induced malignancy.

## Materials and Methods

### *Opisthorchis viverrini* parasite preparation

Metacercariae (MC) of *O. viverrini* were isolated from the naturally infected cyprinid fish by pepsin digestion as previously described (*21*). In brief, fishes were minced with an electric blender, and then minced tissues were digested with 0.25% porcine pepsin, 1.5% HCl in 150 mM NaCl at 37°C in for 2 hours. After tissue digestion, the tissue mixture was filtered sequentially through 1,100, 350, 250, and 140 µm diameter pore size sieves; the final filtrate sedimented by gravity and the aqueous supernatant was discarded. The *O. viverrini* MC enriched-sediment was washed once with 150 mM NaCl (normal saline solution, NSS), and the identity of *O. viverrini* MC confirmed using a stereomicroscope. The *O. viverrini* MCs were stored in NSS at 4°C until used. The newly excysted-juvenile (NEJs) were prepared from MC in 0.25% trypsin in 1×PBS containing 2× 200U/ml penicillin, 200 μg/ml streptomycin (Gibco) (2× Pen/Strep) for 5 min at RT prior to separate the cyst walls from juvenile parasite by insulin needle (*7, 22*). NEJ were transferred to RPMI medium containing 1% glucose, 2 g/L NaHCO_3_, 2× Pen/Strep and 1 µM E-64 protease inhibitor (Thermo Fisher Scientific) and maintained at 37°C, 5% CO_2_ for 60 min before use.

To obtain adult developmental stages of the liver fluke, Syrian golden hamsters (*Mesocricetus auratus*) were infected at 6-8 weeks of age with 50 MC per hamster by intragastric tube (*23*). The hamsters were maintained at the Animal Facility of Faculty of Medicine, Khon Kaen University, Khon Kaen, Thailand. Sixty days after infection, hamsters were euthanized and the liver flukes collected, as described (*22, 23*). The study protocol was reviewed and approved by the Animal Ethics Committee of Khon Kaen University. The study adhered to standard guidelines of the Ethics of Animal Experimentation of the National Research Council of Thailand (approval number ACUC-KKU-61/60).

### **Vector and guide RNA targeting exon 1 of *Ov-grn-1***

To edit the gene *Ov-grn-1* that encodes *O. viverrini* granulin-1 (6287 bp, mRNA GenBank accession FJ436341.1), CRISPR online tools including CRISPR design (http://crispr.mit.edu/) and ChopChop (http://chopchop.cbu.uib.no/) were employed to design a guide RNA (gRNA) targeting exon 1 *Ov-grn-1* gene at nucleotide position 1589-1608, GATTCATCTACAAGTGTTGA (Fig. 1a and 1b). A CRISPR/Cas9 encoding vector encoding the above gRNA under the control of the mammalian U6 promoter and encoding Cas9 (with nuclear localization signal 1 and 2) driven by the CMV promoter was assembled (GeneArt CRISPR Nuclease Vector Kit, Thermo Fisher), and termed pCas-*Ov*-*grn-1* (Supplementary Fig. 1a). *Escherichia coli* TOP-10 competent cells were transformed with pCas-*Ov*-*grn-1* and vector plasmid recovered from cultures of a positive clone (NucleoBond Xtra Midi, Macherey-Nagel GmbH, Germany). The nucleotide sequence of pCas-*Ov-grn-1* was confirmed by Sanger direct sequencing.

### Transfection of liver flukes with pCas-*Ov*-*grn-1*

Twenty mature flukes were transfected with 10 µg pCas-*Ov*-*grn-1* pDNA in ∼500 µl RPMI-1640 (Sigma) by electroporation. The electroporation was performed in 4 mm cuvettes (Bio-Rad) with a single square wave pulse of 125 volts for 20 ms using a Gene Pulser Xcell (Bio-Rad). Flukes were then washed several times with NSS and an additional 5 times with RPMI-1640 containing 2× Pen/Strep. Flukes were cultured in RPMI-1640 containing 2× Pen/Step at 37°C in 5% CO_2_ atmosphere (*7, 24*). Two control groups were included: wild type (WT) mature flukes and ‘mock’ control flukes which were exposed to identical electroporation conditions with RPMI-1640 and 1× Pen/Strep in the absence of plasmid DNA. The adult flukes were observed and collected after 1, 2, 3, 5, 7, 14 and 21 days of culture following pCas-*Ov-grn-1* transfection. RNA and protein was extracted from flukes and *Ov-grn-1* mRNA expression was assessed by RT-qPCR and *Ov*-GRN-1 protein expression was assessed by western blot. Mutations and/or insertions-deletions (INDELs) resulting from CRISPR/Cas were analyzed by CRISPR efficiency estimation (*17, 25*) and MiSeq Next Generation Sequencing (NGS).

MC and NEJ (750 parasites per cuvette) were subjected to square wave electroporation in the presence of pCas-*Ov*-*grn-1* pDNA as described above for adult flukes. The parasites were washed as above and cultured in RPMI complete medium (10% FBS, 2× Pen/Strep) at 37°C in 5% CO_2_ in air. The parasites were collected on days 1, 2, 3 and 5 after transfection and *Ov*-*grn-1* transcript levels were ascertained by RT-qPCR, as above.

### Extraction of nucleic acids

RNA was extracted from pooled or individual flukes using TRIZOL (Invitrogen) according to the manufacturer’s recommendations. Concentration of RNA was estimated by 260 nm by NanoVue spectrophotometer. Genomic DNA from individual or pooled (25 worms) flukes was extracted using the QIAamp DNA Mini Kit (Qiagen, Valencia, CA). The dual RNA and DNA extraction technique was used for some experiments with individual worms using RNAzol®RT (Molecular Research Center, Inc.) and DNAzol (Molecular Research Center, Inc.) (*26, 27*).In brief, each worm was homogenized in RNAzol®RT using a motorized pestle, the DNA and protein from the lysate was precipitated using DNAse-RNAse-free water. The aqueous top solution was transferred into isopropanol to precipitate the RNA. The DNA/protein pellet was resuspended in DNAzol®RT, and DNA extracted as per the manufacturer’s instructions. Individual RNA samples were evaluated for *Ov-grn-1* expression. To assess variations in *Ov-grn-1* transcript levels, individual flukes were assigned into 3 groups; <10% of *Ov-grn-1* transcript fold change (fc) compared to WT, 10-70% *Ov-grn-1* transcript fc compared to WT, and 100-120% *Ov-grn-1* transcript fc compared to WT. Genomic DNA from individual worms was used to assess CRISPR efficiency and estimate mutation levels (*25, 28*). Estimates of mutation efficiency positively correlated with transcripts levels of Ov-*grn-1* transcript level (Fig. 4f). Pooled genomic DNA samples from low to high *Ov-grn-1* transcript levels and high to low percent mutations from experimental groups were prepared for next-generation Illumina sequencing.

### Quantitative Real-time PCR

Complementary DNA (cDNA) was synthesized from parasite total RNA using an iScript cDNA synthesis kit (Thermo Fisher Scientific) prior to proceeding with quantitative real-time PCR (RT-qPCR). RT-qPCR was performed with biological triplicate samples using a SYBR Green kit (TAKARA Perfect Real-time Kit) according to the manufacturer’s recommendation in a Light Cycler 480 II thermal cycler (Roche). Each RT-qPCR reaction consisted of 7.5 µl SYBR Green Master Mix, 0.5 µl (10 µM) each of specific forward and reverse primers for *Ov-grn-1* (Fig. 1b) (forward primer, *Ov*-*grn-1*-RT-F: 5′-GGGATCGGTTAGTCTAATCTCC and reverse primer, *Ov*-*grn-1*-RT-R: 5′-GATCATGGGGGTTCACTGTC), amplifying 359 base pairs (bp) of the product (position 7-365 nt of *O. viverrini* granulin-1 mRNA GenBank accession FJ436341.1), 2 µl of cDNA and distilled water to a final volume of 15 µl. The thermal cycle was a single initiation cycle at 95°C for 3 min followed by 40 cycles of denaturation at 95°C for 30 sec, annealing at 55°C for 30 sec, extension at 72°C for 45 sec and a final extension at 72°C for 10 min. The endogenous actin gene (GenBank accession AY005475) was used as a housekeeping control (*7, 24*) (forward primer, *Ov*-actin-F: 5′-AGCCAACCGAGAGAAGATGA and reverse primer *Ov*-actin-R: 5′-ACCTGACCATCAGGCAGTTC). The *Ov*-*grn-1* transcript fold change was calculated by 2^(-ΔΔCt) method using *Ov-actin* for normalization (*7, 24, 29*). Means and standard deviations were calculated using Graph Pad Prism software; two-way ANOVA.

### Rabbit anti-*Ov-*GRN-1 antiserum and immunoblot analysis

One milligram of adjuvanted recombinant *Ov-*GRN-1 protein (*30*) was used to subcutaneously inject an outbred New Zealand White rabbit. The rabbit was boosted twice with 500 µg of adjuvanted protein and two weeks after the last booster the rabbit was sacrificed for blood collection via cardiac puncture. The Animal Ethics Committee of Khon Kaen University approved the protocols used for animal experimentation, based on the guidelines of the National Research Council of Thailand for Ethics of Animal Experimentation (ACUC-KKU-61/60). *Ov-* GRN-1 protein levels were determined by western blot using rabbit anti-recombinant *Ov*-GRN-1 antiserum. The adult flukes from either WT or Δ*Ov-grn-1* groups were collected individually at days 1, 2, 3, 5, 7, 14 and 21 after electroporation (3 flukes per group). Groups of 3 flukes were homogenized by sonication (Sonics & Materials) in 1× PBS with alternating pulses of 5 sec duration (with 5 sec pause between pulses) for 45 sec at 4°C. The homogenate was centrifuged at 13,000 *g* at 4°C for 30 min and the supernatant collected and stored at −20°C. Protein concentration of fluke homogenates was measured using the Bradford assay and homnogenates were electrophoresed on 15% SDS-PAGE gels. Proteins were transblotted onto nitrocellulose membrane using a Mini Trans-Blot Cell (Bio-Rad). Membrane strips containing 2 µg of total protein were washed with 0.5% Tween-20 in PBS (PBST) then blocked for 1 hour with 5% skimmed milk in PBST. Strips were incubated with rabbit anti-*Ov*-GRN-1 serum or pre-immunization serum diluted 1:50 with 1% skimmed milk in PBST and incubated with shaking for 2 h. Strips were washed then incubated for 1 hour with horseradish peroxidase (HRP)-goat anti-rabbit IgG (Invitrogen) (diluted 1:1,000 in antibody buffer). The strips were washed again and color reactions were detected by enhanced chemiluminescence (ECL) substrate (GE Healthcare Life Sciences) and imaged using an Image Quant LAS 4000 mini (GE Healthcare Life Sciences). As a control protein also derived from the tegument of *O. viverrini* flukes, we assessed the protein expression levels of *Ov-*TSP-2 by western blot using a specific antibody raised to the recombinant protein (*31*). Relative protein expression levels from western blots were measured by densitometry using Image J (https://imagej.nih.gov/ij/download.html). Protein expression levels were compared statistically between groups using independent *t-*tests.

### CRISPR/Cas efficiency and mutation levels estimated by quantitative SYBR green PCR

Adult flukes were collected on days 1, 2, 3, 5, 7, 14 and 21 after pCas-*Ov*-*grn-1* transfection. Each individual fluke DNA was investigated for mutation(s) around the expected double stranded break (DSB) site by using three-primers; 2 forward primers and 1 reverse primer; *Ov-grn-1*-OUT-F, *Ov-grn-1*-OVR-F and *Ov-grn-1*-reverse, respectively. The primer pair of *Ov-grn-1*-OUT-F and *Ov-grn-1*-reverse was used for amplify the fragment flanking (1496-2312 nt) the DSB, while another primer pair (*Ov-grn-1*-OVR-F and *Ov-grn-1*-reverse) was amplify overlap of DSB site (1599-2312) (Fig. 1b). Both primer pairs exhibited equivalent amplification efficiency with genomic DNA form WT flukes. The OUT and OVR amplicons were 450 and 347 bp, respectively, using PCR conditions using 7.5 µl of SYBR Green Master Mix (TAKARA Perfect Real-time Kit), 0.5 µl (0.4 µM) of each primer, 10 ng/µl of gDNA and distilled water to 15 µl. The thermal cycles included initiation for one cycle at 95°C for 3 min followed by 40 cycles of denaturation at 95°C for 30 s, annealing at 55°C for 30 s, extension at 72°C for 45 s, and a final extension at 72°C for 10 min. The SYBR green signal was read every annealing cycle and reported as threshold cycle (Ct). Efficiency of programmed CRISPR/Cas editing was estimated as the ratio of Ct_OUT_:Ct_OVR_ from experimental (A) compared with Ct_OUT_:Ct_OVR_ of control group (B) as described (*17*). The Ct_OUT_:Ct_OVR_ ratio from control group would equal ‘1’ (CRISPR efficiency = 0) since differences were not seen in Ct value from OUT and OVR primers. By contrast, the OVR primer is anticipated to be less efficient than the OUT primer for the experimental group, and hence the Ct_OUT_:Ct_OVR_ is less than ‘1’. Here, we calculated % mutation indirectly by subtraction of the CRISPR/Cas9 efficiency value from ‘1’, as indicated(*17, 32*).

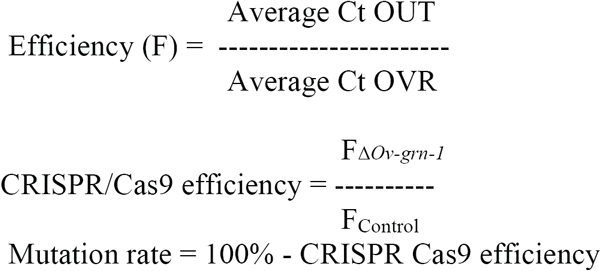

### Illumina-based deep sequencing

The MiSeq NGS library was constructed from pooled DNA samples of *Ov-grn-1* gene-edited adult liver flukes at days 14 and 21 post-transfection. The 173 bp amplicon flanking the DSB was amplified from MiSeq-F (position 1496-1514 nt): 5’ TTCGAGATTCGGTCAGCCG-3’ and MiSeq-R (position 1649-1668 nt): 5’GCACCAACTCGCAACTTACA-3’ primers (Fig. 1b). The amplicon was purified (Agencourt AMPure XP beads, Beckman) and ligated with Gene Read Adaptors Set A (Qiagen) and Illumina compatible adaptor(s) and barcode(s) using QIAseq 1-step Amplicon library kit (Qiagen). The library was quantified using a GeneRead Library Quant Kit (Qiagen) with Illumina index/barcode specific primers included in the kit. The concentration of the MiSeq library was read against known standard libraries provided with the Library Quant Kit. NGS was performed at GENEWIZ, South Plainland, NJ). The MiSeq NGS reads were trimmed of index/adaptor and primer out sequences prior to further analysis for programmed mutations by SnapGene software (GSL Biotech LLC) and CRISPResso analysis platform (*18, 19*) against the target amplicon of the reference (WT) *Ov-grn-1* gene.

### Illumina-based deep sequencing

The MiSeq NGS library was constructed from pooled DNA samples of *Ov-grn-1* gene-edited adult liver flukes at days 14 and 21 post-transfection. The 173 bp amplicon flanking the DSB was amplified from *Ov-grn-1* MiSeq-F (position 1496-1514 nt): 5’ TTCGAGATTCGGTCAGCCG-3’ and *Ov-grn-1* MiSeq-R (position 1649-1668 nt): 5’GCACCAACTCGCAACTTACA-3’ primers (Fig. 1b). The amplicon was purified using Agencourt AMPure XP beads (Beckman) and ligated with index/barcoded adapters compatible with the Illumina system using Gene Read Adaptors Set A with the QIAseq 1-step Amplicon library kit (Qiagen). The library was quantified using a GeneRead Library Quant Kit (Qiagen) with Illumina index/barcode specific primers supplied in the kit. The concentration of the MiSeq library was compared with known standard libraries provided with the Library Quant Kit. The MiSeq reads (NGS performed by GENEWIZ, South Plainfield, NJ) were trimmed of index/adaptor and primer out sequences before downstream analysis for programmed mutations by SnapGene software (GSL Biotech LLC) and the CRISPResso platform (*18, 19*) against the target amplicon from the reference WT *Ov-grn-1* gene.

### Cell proliferation and *in vitro* wound healing assay

To evaluate the effect of *Ov-grn-1* gene editing on liver fluke-driven proliferation of human cholangiocytes, motile WT or Δ*Ov-grn-1* adult flukes were co-cultured with cell of the human cholangiocyte cell line H69 in 24-well Trans-well plates (3 wells per groups) (*7*) containing a 4µm pore size membrane separating the upper and lower chambers (Corning).In brief, 15,000 H69 cells were seeded into the lower chamber of the plate and cultured with complete medium containing DMEM/F12 supplemented with 1× antibiotic, 10% fetal bovine serum, 25 µg/ml adenine, 5 µg/ml insulin, 1 µg/ml epinephrine, 8.3 µg/ml holo-transferrin, 0.62 µg/ml hydrocortisone, 1.36 µg/ml T3 and 10 ng/ml EGF (*33*) for 24 hours, after which the cells were fasted for 4-6 hours in medium supplemented with only one twentieth of the growth factor content of complete medium. Five viable *O. viverrini* adult flukes that had been transfected (or not) with CRISPR/Cas9 *Ov-grn-1* plasmid in a total of 500 µl of RPMI (or medium alone) were placed into the upper chamber of each well. The number of cells in each well was determined at days 1, 2 and 3 using 1× PrestoBlue cell viability reagent (Invitrogen) (*34*) and added to cells at 37°C for up to 1 hour. Cell number was determined at 570 nm and calculated from a standard curve before transforming into relative growth compared to control groups. Cell proliferation assay was carried out in triplicate.

To assess the effect of *Ov-grn-1* knockout on *in vitro* wound healing, 3×10^5^ H69 cholangiocytes in monolayer were grown in 6-well Trans-well plates with a 4 µm pore size. H69 cells were cultured in complete media for 2 days at 37°C then transferred to incomplete media overnight. Monolayers in each well were scratched using a sterile 200 µl autopipette tip (*7, 8, 35*) and washed with PBS twice to remove disconnected cells or debris. Ten transfected adult or control flukes were added to the upper chamber of the Trans-well plate containing the wounded cell monolayer in the lower chamber. The migration rate of cell wound closure was measured at 0, 12, 24 and 36 hours, respectively. Trans-well plates were imaged using an inverted microscope (Nikon) and images of all groups were captured at all-time points quantitatively using Adobe Photoshop CS6. The distances between different sides of the scratch were measured by drawing a line in the middle of the scratch on the captured image (*7, 8, 35, 36*). The analysis of wound healing was carried three times.

### Infection of hamsters with *Ov-grn-1* gene-edited NEJs and assessment of hepatobiliary histopathology

Thirty male Syrian golden hamsters, 6-8 weeks of age, were obtained from the Animal Unit, Faculties of Medicine, Khon Kaen University (approval number ACUC-KKU-61/60). The hamsters were randomly divided into three groups of 10 per group: uninfected control, infected with WT flukes and infected with Δ*Ov-grn-1* flukes. Each hamster was infected with 100 active NEJs through intragastric intubation; the uninfected control group was fed normal saline solution instead of NEJ (*23*). Hamsters (5 animals per cage) were contained under conventional conditions and fed a stock diet (C.P. Ltd., Thailand) and water *ad libitum* until they were euthanized (*23*). Following euthanasia, five hamsters from each group were necropsied for histopathological assessment of the hepatobiliary tract at day 14 and at day 60 post-infection (*23*). The hamsters were euthanized by overdose of anesthesia with diethyl ether. Subsequently, blood was obtained by cardiac puncture and the livers were removed. Fluke numbers were counted from two livers of both, WT and Δ*Ov-grn-1* groups at day 60 after infection of hamsters and compared with unpaired two-tailed *t*-test. The left and right lobes of the liver from five hamsters were dissected, cross-sectioned and each lobe was divided into three parts. The liver fragments were fixed in 10% buffered formalin and stored overnight at 4°C before processing. Formalin-fixed liver was dehydrated through an ethanol series (70, 95 and 100%), cleared in xylene and embedded in paraffin. Paraffin embedded sections of 4 µm thickness, cut by microtome, were stained with hematoxylin and eosin (H&E) or Picro-Sirius Red, or probed with anti-ACTA2 antibodies, and analyzed for pathologic changes (below).

### Biliary hyperplasia

H&E staining was used to assess pathological changes. The sections were deparaffinized in 100% xylene, rehydrated with descending series of alcohol, stained with H&E for 5 min, dehydrated with an ascending series of alcohol, cleared with 100% xylene, mounted on a slide with permount media, and slides were dried at 37°C overnight and photographed under light microscopy. Images (200×) from H&E stained liver sections from 5 untreated control hamsters or 5 hamsters per treatment group (WT and Δ*Ov-grn-1* groups) were assessed. Thickness (width) of the bile duct epithelium from each thin liver section was measured with ImageJ at eight equidistant positions around the bile duct. To compensate for outliers the median width for each bile duct was used for graphing and analysis. The two-way ANOVA Holm-Sidak multiple comparison test was used to compare each group at each time point.

### Fibrosis

Two stains were used separately to assess biliary fibrosis. First, sections were stained with Picro-Sirius Red (Abcam, Cambridge Science Park, UK). Sufficient Picro-Sirius Red solution was applied to completely cover the tissue sections on the slide, the stained slide was incubated at ambient temperature for 60 min, rinsed in two changes of acetic acid solution, and dehydrated through two changes of absolute ethanol. Slides were cleared with 100% xylene, mounted in Permount, dried at 37°C overnight, and examined and photographed by light microscopy to document collagen surrounding the bile ducts. ImageJ was used to auto-color balance the images using the macro written by Vytas Bindokas; Oct 2006, Univ. of Chicago (https://digital.bsd.uchicago.edu/docs/imagej_macros/_graybalancetoROI.txt) followed by application of the MRI fibrosis tool to determine percentage area of red-stained fibrosis at default settings (red 1: 0.148, green 1: 0.772, blue 1: 0.618, red 2: 0.462, green 2: 0.602, blue 2: 0.651, red 3: 0.187, green 3: 0.523, blue 3: 0.831) (*37*). Twenty discrete images (200×) stained with Picro-Sirius Red from each animal (five hamsters per treatment group) were assessed (in total, 100 images per group). Kruskal-Wallis with Dunn’s multiple comparison test was used to compare the findings due to the range of data points among the groups.

Fibrosis was also assessed via levels of smooth muscle alpha-actin (ACTA2). Liver sections from hamsters at day 60 post-infection were deparaffinized three times with 100% xylene, 5 min each. Sections were rehydrated with ascending series of ethanol; 100%, 3 times, 3 min each, 95% 3 times, 3 min each, 70% for 3 min, followed by thorough washing in tap water for 5 min, distilled water for 5 min, and PBS for 5 min. Thereafter, slides were incubated in citrate buffer pH 6.0 (citric acid (anhydrous) 1.92 g and Tween 20 0.5 ml in total volume of DW 1000 ml) at 110°C for 5 min, allowed to cool for 20 min, and then washed in PBS 3 times, 5 min each. Sections were then blocked with 5% bovine serum albumen (BSA) for 30 min in a humidified chamber and washed in PBS 3 times, 3 min each with occasional shaking. The slides were probed with Alexa Fluor 594 –labeled anti-ACTA2 antibody (Abcam) diluted 1:200 in 1% BSA in PBST, 18 hours at 4°C in a humidified atmosphere. Lastly, slides were washed in PBS 3 times, 3 min each with occasional shaking, mounted in glycerol diluted 1:4 with PBS and examined under bright and fluorescence light (Zeiss Axio Observer, with AxioVision SE64 Rel. 4.9.1 software, Jena, Germany). Images with a bile duct containing a fluke were selected and ImageJ was used to manually select a narrow strip surrounding the bile duct epithelial layer that excluded potential blood vessels (any enclosed curved oval-like shape). Three liver regions not including the bile duct or blood vessels were manually selected for background fluorescence measurements, each comprising 5-10% of the image. The fluorescence intensity of the bile duct epithelial strip was measured and blanked against the average of the three background regions and reported as average intensity per cm^2^ at 300 PPI (pixels per inch). A total of 25-30 distinct bile duct images per group (3 animals) were assessed. Zero values from the uninfected group were deemed to have a value of 1 to allow a log axis. The groups were compared with one-way ANOVA with Holm-Sidak multiple comparison test.

## ACKNOWLEDGMENTS

We thank Suwit Balthaisong for technical assistance. This study was supported by the Thailand Research Fund through the Royal Golden Jubilee PhD Program (grant no. PHD/0111/2557 to PA and TL), the National Cancer Institute (NCI), US National Institutes of Health (NIH) award R01CA164719 (AL, TL, PJB,) and the National Health and Medical Research Council (NHMRC award APP1085309 to AL, JS and TL, and senior principal research fellowship APP1117504 to AL). The content is solely the responsibility of the authors and does not necessarily represent the official views of the NIAID, NCI, NIH or NHMRC. These studies were supported in part by Wellcome Trust Strategic Award number 107475/Z/15/Z (Karl F. Hoffmann, principal investigator; PJB, co-investigator).

## AUTHOR CONTRIBUTIONS

T. Laha, P.J. Brindley, W. Ittiprasert, P. Arunsan, N. D. Young, A. Loukas, J. Sotillo, M. Smout, S. Chaiyadet, C. Cochran, and V. Mann conceived and designed the research; P. Arunsan, M. Smout, J. Sotillo, S. Karinshak and W. Ittiprasert performed the research; P. Arunsan and M. Smout recorded the findings; T. Laha, P.J. Brindley, M. Smout and W. Ittiprasert contributed reagents and analytic tools; T. Laha, P.J. Brindley, M. Smout, J. Sotillo and P. Arunsan completed the experiments; T. Laha, P. Arunsan, W. Ittiprasert, J. Sotillo and M. Smout analyzed data; T. Laha, P. Arunsan, W. Ittiprasert, M. Smout, A. Loukas and P.J. Brindley prepared figures and wrote the paper.

The authors declare no competing interests.

## Supporting information

**Supplementary Figure 1.**
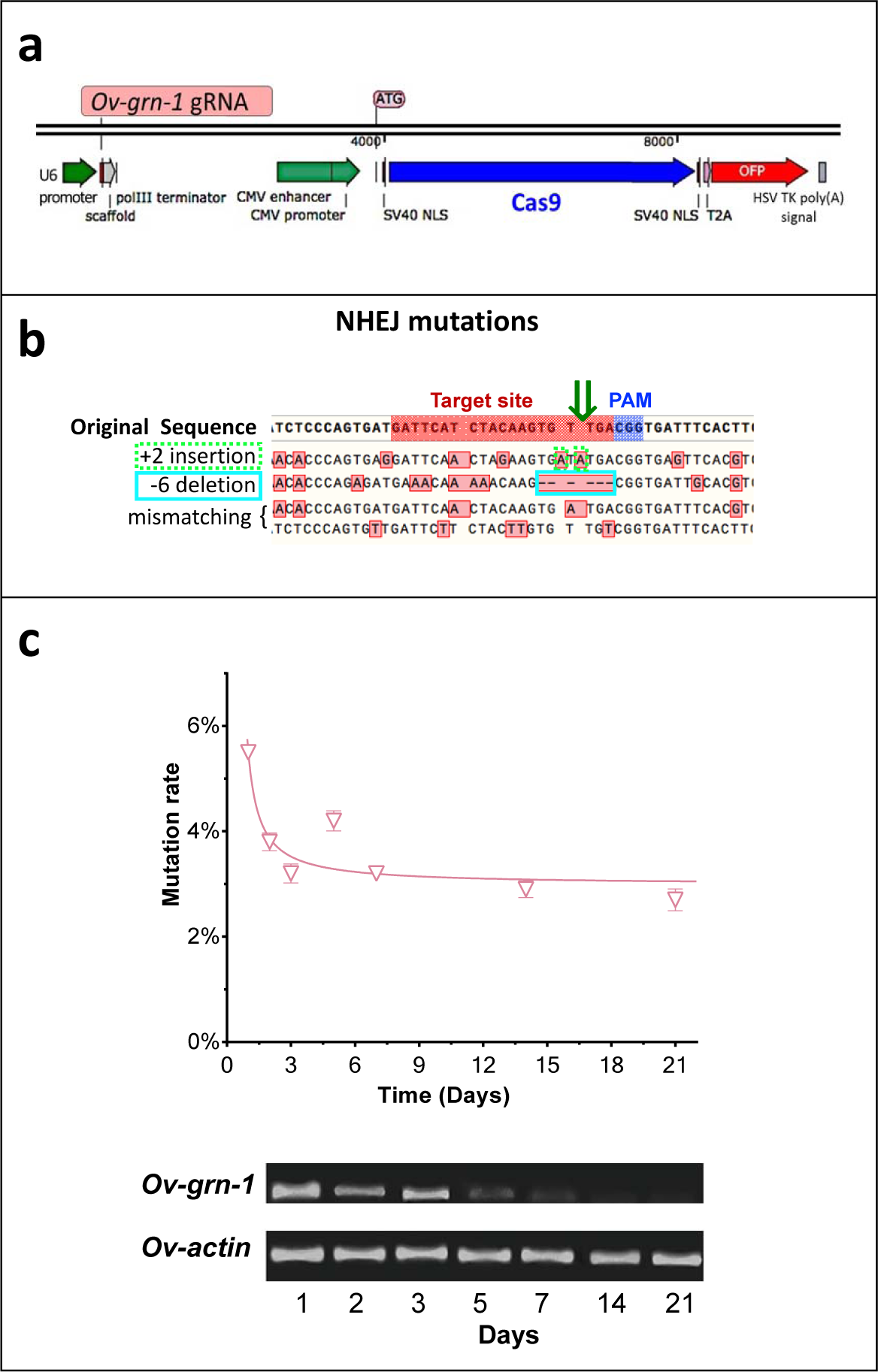
CRISPR/Cas9 targeting *Ov-grn-1* design and flukes transfection. **a**, *Ov-grn-1* guide RNA insertion and Cas9 cassette. **b**, The original *Ov-grn-1* sequence with the target (red text) and PAM (blue text) sites highlighted. The dark green arrow indicates the specific CRISPR target site. Deep sequencing is used to determine specific mutations and the four sequences below the original sequence show examples of successfully mutated *Ov-grn-1* genes with variations with red boxed bases highlighted mutations induced by the CRISPR/Cas9 transfection. All four examples show a range of insertion, deletion or mismatch mutations boxed in red. The “+2 insertion” with a dotted light green box highlights two alanine insertions near the target site. The “-6 deletion” highlights six absent base pairs with a light blue box which include the target site. **c**, The mutation rates were calculated at each time point from Δ*Ov-grn-1 in vitro* cultured adult flukes with qPCR and the averages and standard deviation were determined. RT-qPCR electrophoresis gel (lower image) shows *Ov-grn-1* RNA levels are reduced in Δ*Ov-grn-1* adult flukes.

**Supplementary Figure 2.**
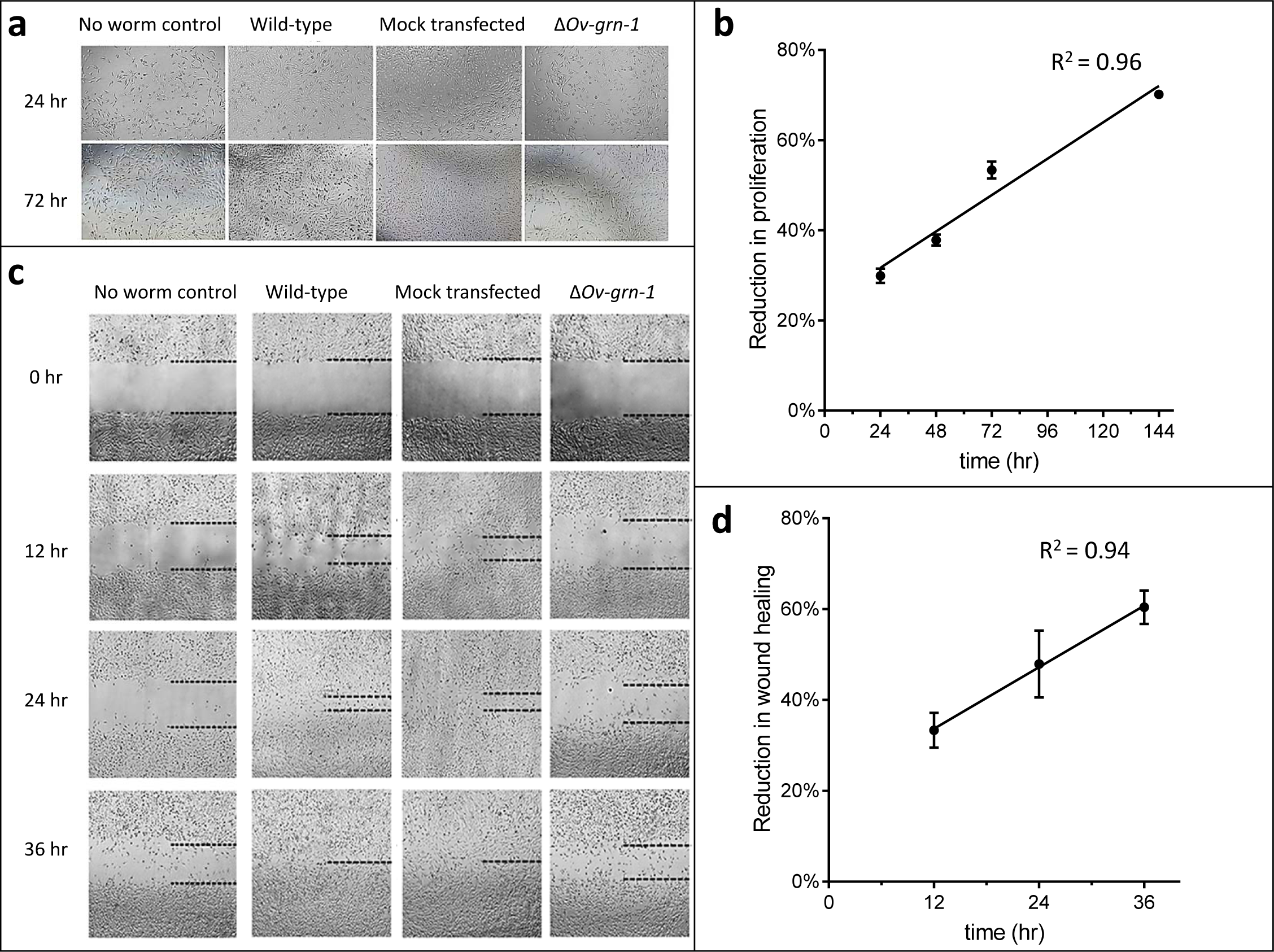
Extended image set showing Δ*Ov-grn-1* adult fluke ES products induce less *in vitro* cell proliferation and wound repair. **a**, Representative cell proliferation images of H69 immortalized human cholangiocyte cell line co-cultured with flukes in Trans-well plates; shown at 24 and 72hrs. **b**, Comparing the difference in cell proliferation values from Fig. 2b between ES products from the *ΔOv-grn-1* flukes and ES from WT flukes revealed a linear correlation of reduction of cell proliferation that increases over time. **c**, Representative image of scratch wound repair of cultured cholangiocytes when H69 cells were co-cultured in Trans-well plates with flukes; time points from 0 - 36 hours post-scratch. Dotted line indicates the margin of the wound. **d**, Scratch wound repair assay quantified values from Fig. 2d showed an increasing linear correlation over time of reduced wound closure when plotting the difference between the Δ*Ov-grn-1* and the WT fluke groups. Panels **b** and **d**: mean ± SD, three biological replicates.

**Supplementary Figure 3.**
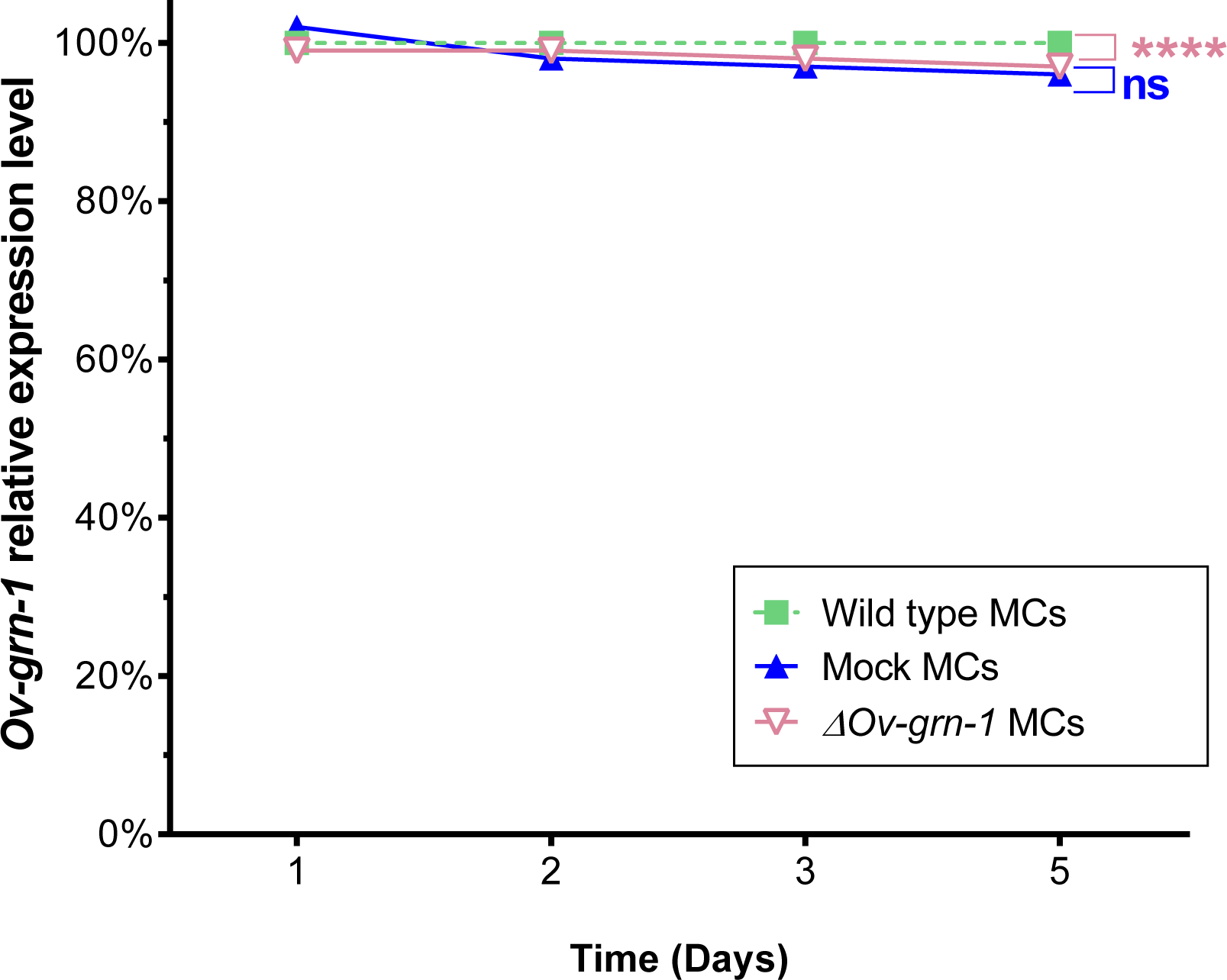
CRISPR/Cas9 ineffective at silencing gene expression in metacercariae (MC). Expression of *Ov-grn-1* mRNA in wild type (WT), mock-transfected, and Δ*Ov-grn-1* MC over five days as quantified with qPCR and plotted relative to the WT untreated group; mean ± SD from three replicates. \*\*\*\**P*<0.0001; Δ*Ov-grn-1* compared to WT control group, *P*>0.05; non-significant (ns) difference seen between mock (control) group using the two-way ANOVA Holm-Sidak multiple comparison test.

**Supplementary Figure 4.**
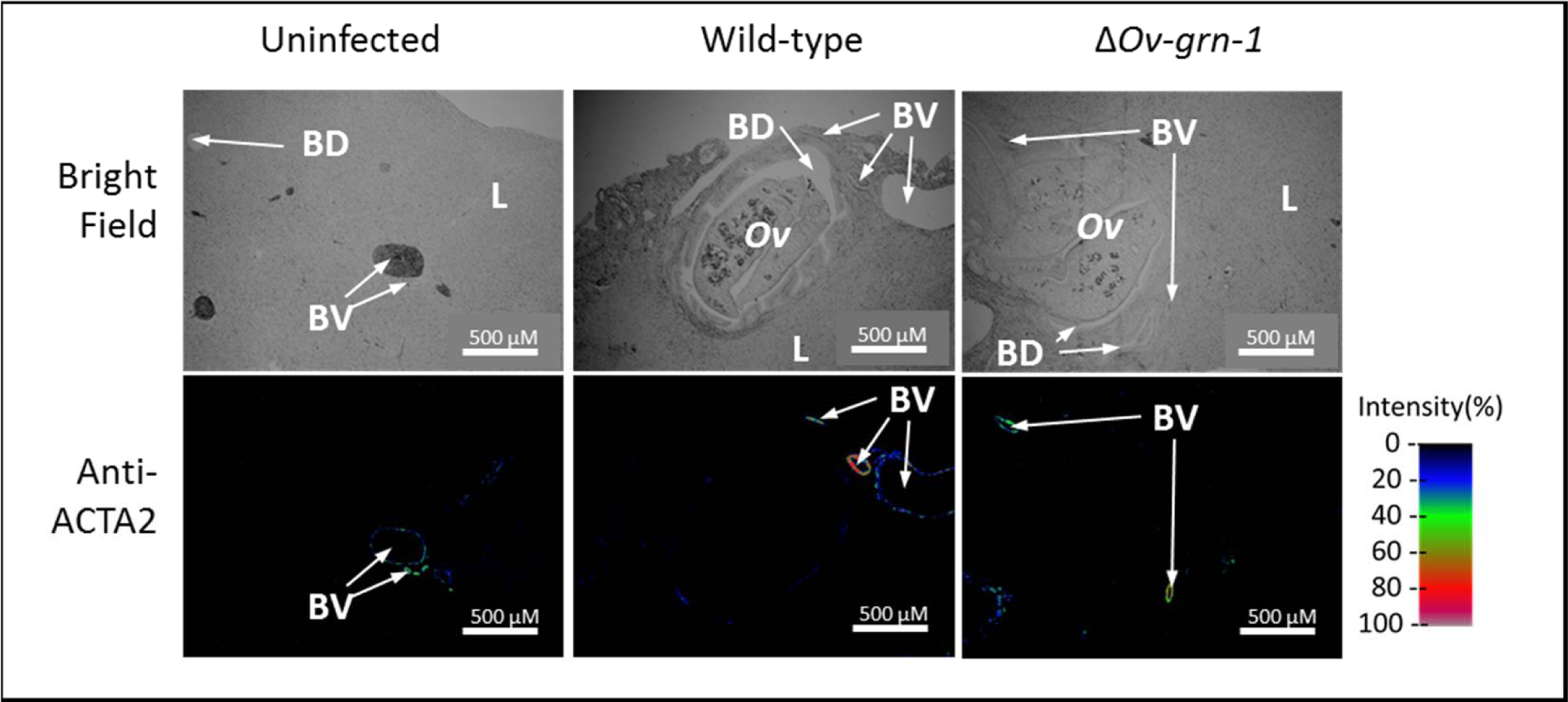
Representative wide-angle view of anti-ACTA2 immunofluorescence and bright field liver sections. Thin liver sections presented in bright field or under fluorescent light following probing with anti-ACTA2 antibody; fluorescence intensity shown on a blue/green/red scale. These images were used to create the bright field/immunofluorescence overlay and zoomed images in Fig. 4d.

